# The lateral habenula is required for maternal behavior in the mouse dam

**DOI:** 10.1101/2024.02.12.577842

**Authors:** Jessie Benedict, Robert H. Cudmore, Diarra Oden, Aleah Spruell, David J. Linden

## Abstract

Mammalian parenting is an unusually demanding commitment. How did evolution co-opt the reward system to ensure parental care? Previous work has implicated the lateral habenula (LHb), an epithalamic nucleus, as a potential intersection of parenting behavior and reward. Here, we examine the role of the LHb in the maternal behavior of naturally parturient mouse dams. We show that kainic acid lesions of the LHb induced a severe maternal neglect phenotype in dams towards their biological pups. Next, we demonstrate that through chronic chemogenetic inactivation of the LHb using DREADDs impaired acquisition and performance of various maternal behaviors, such as pup retrieval and nesting. We present a random intercepts model suggesting LHb-inactivation prevents the acquisition of the novel pup retrieval maternal behavior and decreases nest building performance, an already-established behavior, in primiparous mouse dams. Lastly, we examine the spatial histology of kainic-acid treated dams with a random intercepts model, which suggests that the role of LHb in maternal behavior may be preferentially localized at the posterior aspect of this structure. Together, these findings serve to establish the LHb as required for maternal behavior in the mouse dam, thereby complementing previous findings implicating the LHb in parental behavior using pup-sensitized virgin female mice.

**Significance Statement:** Work conducted using rats in the 1990s suggested an important role for the LHb in maternal behavior, but this area of research has since lain dormant. In the interim, the LHb has been garnering attention as a hub for punishment signaling. Recently, interest in the LHb’s role in maternal behavior was renewed, with an important paper examining LHb function during pup-directed behaviors in pup-sensitized virgin female mice. But it is unknown how closely pup-directed behaviors in sensitized virgin females may mimic maternal behavior in natural mouse mothers. This work demonstrates the importance of the LHb in the regulation of natural maternal behavior in the mouse.

## Introduction

Reproduction largely employs one of two strategies: 1) engage in a massive initial energy expenditure, laying a clutch of 7,500 eggs every year like the Atlantic salmon, and then leaving offspring survival to chance. Or 2) gestate a limited number of offspring, like the mouse, investing in their care until they become independent. The second model, favored by birds and mammals, imposes a significant cost on the parent. Parenting involves making repeated costly choices — limiting foraging time to defend the nest, sharing food, and confronting predators rather than hiding or escaping. Given these costs, the neural circuitry coordinating the shift from self-prioritization to offspring-prioritization should be tightly controlled and robustly rewarding. In addition, maternal behavior is a complex array of behaviors necessitating specific adjustment across offspring development. For instance, mouse pups old enough to locomote independently no longer require maternal retrieval but instead require new maternal vigilance to ensure pups do not wander dangerously far from the nest. Thus, parenting behavior requires robust reward signaling and flexible decision-making.

Mounting evidence implicates the lateral habenula (LHb), as a neuroanatomical intersection of parenting behavior, flexible decision-making, and reward signaling. LHb lesions prevent the onset of maternal behavior in the rat (Corodimas et al., 1992, 1993; Matthews-Felton et al., 1995; Felton et al., 1998, 1999). The LHb is also required for flexible non-maternal behaviors, with LHb inactivation causing task-switching and perseverative errors in the rat (Baker et al., 2015, 2017). LHb neurons mediate approach of appetitive stimuli and avoidance of aversive stimuli in the mouse (Stamatakis and Stuber, 2012; Proulx et al., 2014; Stephenson-Jones et al., 2020; Lalive et al., 2022; Mondoloni et al., 2022). The anticipation of punishment and reward omission both cause phasic increases in LHb activity in rhesus macaques (Matsumoto and Hikosaka, 2007). By virtue of its bidirectional connectivity with both the ventral tegmental area (VTA)/substantia nigra pars compacta (SNc) and the raphe nuclei, the LHb serves as an important integration site for the dopamine and serotonin systems in the brain (Reisine et al., 1982; Amat et al., 2001; Yang et al., 2008; Kim, 2009; Omelchenko et al., 2009; Jhou et al., 2013; Stamatakis et al., 2013; Amo et al., 2014; Brown et al., 2017). Unsurprisingly, the LHb has been implicated as an important node in the pathophysiology of depression in rodents, rhesus macaques, and humans (Matsumoto and Hikosaka, 2007; Sartorius and Henn, 2007; Yang et al., 2008, 2018; Li et al., 2011; Proulx et al., 2014; Shabel et al., 2014), making the LHb an exciting area of research for peripartum depression and anxiety, which affects 13-19% of human mothers (O’Hara and McCabe, 2013; Dennis et al., 2017; Shovers et al., 2021; Wang et al., 2021).

Recently, interest in the LHb’s role in parenting behavior has been renewed (Hu et al., 2020; Li, 2022; Mondoloni et al., 2022; Lecca et al., 2023) with an exciting paper by Lecca et al. (2023), examining LHb function during pup-directed behaviors in sensitized virgin female mice. Sensitized virgin female mice are virgin female mice who, after repeated exposure to pups, begin to perform alloparental behaviors. Sensitized virgin females are often used in place of naturally parturient mouse dams, likely to improve experimental throughput. But it remains unclear how the neural dynamics governing pup-directed behaviors in sensitized virgin females compare to those governing maternal behavior in natural dams. Virgin mice learn alloparental care best through social transmission from an experienced dam (Carcea et al., 2021). This is a notably different mechanism than naturally parturient maternal behavior. Though sensitized virgin female mice can perform on par with dams in some pup-directed behaviors, such as latency to retrieve a pup to the nest (Stolzenberg and Rissman, 2011), when motivation is specifically assayed, stark differences between virgins and dams emerge (Gandelman et al., 1970; Salais-López et al., 2021). In a retrieval assay where sensitized virgin female mice and dams could retrieve a live pup or dead pup from each arm of the T-maze, the sensitized virgin female group retrieved live pups a total of two times. In the same period, the dam group retrieved dead pups 83 times, and live pups 83 times (Gandelman et al., 1970). Thus, sensitized virgin mice are protected from over-investment in others’ offspring even as dams have undergone such a robust shift to offspring-prioritization that they continue to perform maternal behavior for dead pups.

Here, we have sought to test the hypothesis that the LHb is necessary for maternal behavior by using primiparous naturally parturient mouse dams and their litters to examine the effect of LHb kainic acid lesions and chemogenetic inactivation using DREADDs on maternal behavior.

## Methods

### Experimental Animals

Pup-naïve virgin female C57BL/6 mice (RRID:IMSR_CRL:027) aged 6-9 weeks were randomly assigned to treatment group for all experiments. Male C57BL/6 mice (RRID:IMSR_CRL:027) were used for breeding purposes only, as all experiments herein examine maternal behavior. All experiments were approved by [redacted for double-blind review]. Animals were housed on a 12:12 light/dark cycle. Food and water were available *ad libitum*.

### Kainic Acid Lesion Experiments

#### Stereotaxic Injection Surgery

Mice were anesthetized using vaporized isoflurane (5% for induction, 1.5% for maintenance) and placed in a stereotactic frame (Leica, Angle 2). Opthalmic ointment was applied and rectal temperature and respiration rate were monitored throughout surgery. The mice were shaved and iodine was swabbed on the scalp. Mice were given lidocaine (0.5% lidocaine, 0.02 ml) and dexamethasone (0.20 mL at 0.4 mg/mL) subcutaneously under the scalp. Mice received diazepam (2.5 mg/kg, intraperitoneal (IP) injection) as an analgesic and an anti-convulsant, and enrofloxacin (2.5 mg/kg IP injection) as an injectable antibacterial solution. An incision was made on the scalp to reveal the skull surface. Bregma and lambda were aligned in the mediolateral and dorsoventral planes within 30 mm of each other, and the points 2 mm lateral of bregma in each hemisphere were aligned within 30 mm of each other in both the anteroposterior and dorsoventral planes. A craniotomy was performed using a rotary tool affixed to the stereotactic frame. The drill rotated a 0.9mm drill burr which was irrigated with ice-cold saline. Injections were performed with a Nanoject II (Drummond) using a glass pipette with a tip diameter of 12.5-22.5 mm. Depending on experimental group assignment, either sterile saline (0.9% Sodium Chloride, Hospira), or kainic acid (1mg/ml dissolved in sterile saline, Tocris; Catalog No: 0222), were injected into the lateral habenula. Injections of 101.2 nL volume were made bilaterally at −1.50 anteroposterior (AP), +/− 0.42 mediolateral (ML), −2.70 dorsoventral (DV). Approximately 5 minutes elapsed before retracting the pipette from the brain. During the retraction, slight negative pressure was applied to the pipette through the Nanoject II. The scalp was sutured with absorbable polyglycolic acid (PGA) suture and iodine was reapplied. Mice were then given IP buprenex (0.05 mg/kg) and 0.5 mL IP sterile saline to prevent dehydration and placed under a heat lamp for observation. Once a mouse awoke from anesthesia and was ambulating normally, it was returned to a shared cage. Mice were supplied with hydration gel postoperatively and their sutures were monitored daily for seven days.

#### Breeding and Pregnancy

Animals were given at least one week to recover postoperatively and were then housed in a harem breeding scheme with 2-3 females per breeding male. Mice remained group housed until ∼E17 when pregnant females were moved to cages within individual behavior boxes to allow for continuous video monitoring as they neared parturition.

#### Naturalistic Maternal Behavior Tests

Lactation day 0 (LD0) was demarcated as the first day live pups were seen in the cage by 1200 (h) hours. Naturalistic maternal behavior tests (MBTs) were conducted on LD1. Standard MBTs were planned to start on LD2, but these were never performed since nearly all the kainic acid-treated dams’ litters had died of neglect by LD2. The naturalistic MBTs were scored from 30-minutes of video in which all dams demonstrated activity, between 1400-1600 h.

#### Behavior Scoring

The 30-minute naturalistic MBT video was scored blind, using custom software detailed in [redacted for double-blind review], consistent with standard MBTs (Numan and Callahan, 1980; Matthews-Felton *et al*., 1995; Felton *et al*., 1998; Nephew and Bridges, 2011; Wu *et al*., 2014). Thirty 10-second video windows were randomly selected from each assay and scored in a randomized order. Each window was scored for the presence or absence of the following behavioral states: all pups gathered, all pups nested, arched-back hovering/visible nursing, time with pups, pup interactions (retrieval, inspection, anogenital licking), nest building, solo activity, solo rest, pup lacking visible milk spot, dead pup visible. Additionally, a blinded experimenter scored each nest at the end of the MBT. Nests were scored using integers 0 to 4 as follows: 0, no nest attempted; 1, poor nest, not all of the nesting material was used, lacks structure; 2: fair nest, all the nesting material was used but the nest lacks structured walls; 3, good nest, all nesting material was used and the nest has low walls; 4, excellent nest, all the nesting material was used and the nest has high structured walls (Numan and Callahan, 1980). Scores for all pups nested were not reported in Extended Figure 1-1 since all values were identical to those reported for all pups gathered.

### Chronic Chemogenetic Inactivation with DREADDS

#### Stereotaxic Injection Surgery

All procedures are identical to those described in the Kainic Acid Lesion Experiments – Stereotaxic Injection Surgery subsection, except for the following differences: Mice received preoperative buprenex (1mg/kg) rather than diazepam. The drill burr for the craniotomoy was 1.2mm rather than 0.9mm, to enable multiple injection sites. Rather than kainic acid or saline, depending on group assignment, either experimental AAV2-hSyn-hM4D(Gi)-mCherry (RRID: Addgene_50475) or control AAV2-hSyn-mCherry virus (RRID: Addgene_114472) was injected into the LHb. Injections of 101.2 nL were made bilaterally at −1.45 AP and −1.55 AP, +/− 0.42 ML, −2.70 DV or, later, at −1.28 AP and −1.38 AP and −1.48 AP, +/− 0.42 ML, −2.70 DV, as immunohistochemical results from early cohorts showed sparser transfection of the anterior-most aspect of the LHb and so coordinates were adjusted.

#### Breeding and Pregnancy

Procedures were identical to those described in Kainic Acid Lesion Experiments – Breeding and Pregnancy.

#### Agonist-21 Administration

Approximately 24 hours before parturition (range 6-40 hours), each mouse began receiving subcutaneous injections of the hM4Di agonist called Agonist-21 (HelloBio, dose: 3 mg/kg, HB4888) every 6 hours (0900, 1500, 2100, 0300 h) to maintain putative neural inactivation in the lateral habenula (Thompson et al., 2018; Jendryka et al., 2019; Ferrari et al., 2022). Once at least six hours had elapsed since parturition, the uterus was palpated and if no additional pups were suspected, injections were shifted to the IP route to minimize discomfort. Mice were weighed at 0900 h to determine their dose for that day.

#### Standard Maternal Behavior Tests

LD0 was demarcated as the first day live pups were seen in the cage by 1200 h. Behavior experiments were conducted on LD1, LD2, LD3, and LD4, during the first hour of the light cycle. Following Agonist-21 administration, dams were placed in a clean separation chamber for 30 minutes. While the agonist took effect, the cage was cleaned, food and water were removed for the duration of the assay, and new nesting material was supplied. Pups were examined for the presence or absence of milk spots, signs of physical injury, and weight gain. Just before the 30-minute mark, pups were returned to the clean cage and scattered in the three corners opposite a new nestlet. The video recording began at the dam’s moment of return to the cage, which commenced the 30-minute MBT.

#### Behavior Scoring

The standard MBT differs from the naturalistic MBTs (see Kainic Acid Lesion Experiments) in that it includes a separation period before the dam is reintroduced to scattered pups. Therefore, the behavior scoring for standard MBTs includes scores for delay to initiate first pup retrieval and delay to complete pup retrieval (Extended Figure 3-1). Since the assay duration was 1,800 seconds (30 minutes), animals that failed to initiate or complete pup retrieval by that point received a score of 1,800 seconds. The standard MBT scores for solo rest, pup lacking visible milk spot, and dead pup visible are not included in Extended Figure 3-1 since all values for both groups were equal to zero.

### All Experiments

#### Perfusion and Tissue Processing

On LD4, dams were briefly anesthetized with isoflurane and injected with a ketamine/xylazine cocktail (4 mL/kg) to achieve deep anesthesia. When unresponsive to a toe pinch, the dams were then myocardially perfused with ice-cold 1X PBS and ice-cold 4% PFA (Electron Microscopy Sciences, 20% solution, EM Grade #15713, diluted to 4% in PBS). Brains were postfixed overnight in 4% PFA and then placed in 15% sucrose until they sunk. The same procedure was then repeated using a 30% sucrose solution. The brains were sectioned coronally on a freezing microtome. 40-micron thick sections were collected, and alternate sections were then used for immunohistochemistry and histology analysis.

#### Immunohistochemistry

Sections were washed 3 times in wash buffer (1x PBS + 0.3% Triton-X in a 24-well plate) for at least 5 minutes per wash. Next, they were placed in blocking serum (wash buffer + 5% Normal Goat Serum (Jackson ImmunoResearch, RRID: AB_2336990) for one hour at room temperature or overnight at 4℃. Primary antibodies, anti-NeuN (Millipore Cat# MAB377, RRID: AB_2298772) and anti-GPR151 (Sigma-Aldrich Cat# SAB4500418, RRID: AB_10743815), were added to fresh blocking serum and sections were incubated for 1-2 hours at room temperature or overnight at 4℃. After this step, sections were washed 3 times in wash buffer for at least 5 minutes per wash. Sections were then incubated with secondary antibodies Alexa Fluor 488-AffiniPure Goat Anti-Mouse IgG (H+L) (Jackson ImmunoResearch Labs Cat# 115-545-003, RRID:AB_2338840) and <u>Alexa Fluor 647-AffiniPure Goat Anti-Rabbit IgG (H+L) (min X Hu,Ms,Rat</u> <u>Sr Prot)</u> (Jackson ImmunoResearch Labs Cat# 111-605-144, RRID: AB 2338078) in fresh blocking serum for 2-4 hours at room temperature or overnight at 4℃. Lastly, sections were washed 3 times in wash buffer for at least 5 minutes per wash before being mounted with ProLong Diamond AntiFade Mountant with DAPI and allowed to dry overnight.

#### Microscopy and Histological Analysis

All microscopy and cell counting was conducted blinded to treatment group using a Zeiss 800 inverted confocal microscope. Tiled z-stacks were collected with a 20x objective to capture the entire LHb, with 8-11 images analyzed per animal. Image files were stitched using ZEN Blue software (RRID: SCR_013672) and then exported for cell and volume quantifications. Imaris software (version 9.8, RRID: SCR_007370) was used to outline the LHb across all z-slices of a given image, guided by a combination of NeuN and Gpr-151 staining. Then, all cells within outlined LHb borders were exhaustively counted, using NeuN immunofluorescence for counting neurons in the kainic acid lesion experiments and mCherry immunofluorescence for counting transfected cells in the chronic chemogenetic inactivation experiments. Separate blinded scoring was performed of off-target transfections in the chemogenetic inactivation experiments. Every brain section was given a subjective 0-3 score (−, +, ++, +++) for each of five neighboring brain regions: the hippocampus, the medial habenula, the paraventricular nucleus of the thalamus, the central lateral nucleus of the thalamus, and the anterior pretectal nucleus. The scoring rubric was: no cells transfected (−), a minority of cells were transfected ∼5-15 (+), a moderate number of cells were transfected, ∼15-100 (++), a large number of cells were transfected, 100+ (+++). Each animal’s off-target transfection score was averaged across each section containing a given off-target region, and these final values were analyzed.

#### Statistical Analysis

R Studio (version 2023.09.0+463) was used with R Project for Statistical Computing (RRID: SCR_001905, version 4.3.1) for analysis. All error bars in the figures show the median and interquartile range (IQR) and all univariate behavior analyses are nonparametric Wilcoxon Rank Sum tests. Pearson correlations were used to compare total neuron counts/total transfected cell counts to behavior measures (Figure 2D, Extended Figure 6-1). Spearman correlation was used to test for correlation between behavior results and non-linear scale histology data from off-target brain region transfections (Extended Figure 6-2). The random intercepts linear mixed models were all conducted using the R package: lme4 (RRID: SCR_015654). All statistical tests, particularly the linear mixed models, were conducted in direct consultation with [redacted for double-blind review]. The random intercepts models were tested for Akaike Information Criterion (AIC) value against multivariate fixed-effect linear models, univariate linear models, and random slope/random intercept and random slope mixed linear models. In all models presented, the random intercept model was the most parsimonious, with the lowest AIC score. All analysis was conducted on a Mac-Mini (3.2 GHz 6-Core Intel Core i7, running macOS Ventura 13.2).

#### Data Availability

All of the data presented in this manuscript is made available in a downloadable folder, along with all of the code for the multilevel multivariate models

## Results

### LHb kainic acid lesions induced a severe maternal neglect phenotype in dams

Seeking to extend prior findings that kainic acid lesions of the lateral habenula prevented the onset of maternal behavior in the rat (Matthews-Felton et al., 1995), we tested the effects of kainic acid LHb lesions in the mouse dam. Pup-naïve virgin female mice received bilateral LHb injections of either saline or kainic acid. Following recovery, breeding, and pregnancy, the mice were moved to individual behavior boxes with continuous video monitoring [citation redacted for double blind review] on E17 to habituate prior to parturition (Figure 1A). While standard maternal behavior tests (MBTs) were planned to begin on lactation day 2 (LD2), half of all dams’ (8/16) litters had died by then. Instead, naturalistic MBTs (Carcea et al., 2021; Dvorkin and Shea, 2022; Schuster et al., 2023) were scored from footage obtained on LD1 in which all dams showed activity. Naturalistic MBT analysis showed that 7/8 kainic acid-treated dams failed to engage in any maternal behavior, resulting in the death of 100% of their litters by LD2. Importantly, dams did not cannibalize their pups, but left them strewn about the cage as if they were part of the environment, occasionally sniffing or stepping on them (Figure 1B; Extended Figure 1-3). The neglected pups exhibited no visible milk spots (Figure 1B; Extended Figure 1-1). Naturalistic MBTs do not begin by manually scattering the pups to specifically test pup retrieval, but pups do intermittently fall out of the nest. The best proxy variable for retrieval behavior is “all pups gathered” (Figure 1C), which shows that 7/8 saline-treated dams had all of their pups gathered in nearly all of the video windows that were scored. Simultaneously, 7/8 of the kainic acid-treated failed to have all their pups gathered in nearly all the scored video windows (*p* >. 01, Wilcoxon rank sum test in all comparisons, unless otherwise stated). Arched-back hovering/visible nursing scores (*p* > .01) (Figure 1D), were similar to “all pups gathered”. Nest scores (Figure 1E) show that most saline-treated dams built excellent nests with high structured walls and used all their available nesting material, whereas kainic acid-treated dams’ nests scored far worse (*p* > .01) (see Methods, Behavior Scoring for more detail on nest scoring). For a table of all behaviors measured, see Extended Figure 1-1. For example video footage of saline-treated and kainic acid-treated dams, see Extended Figures 1-2, 1-3.

**Figure 1.**
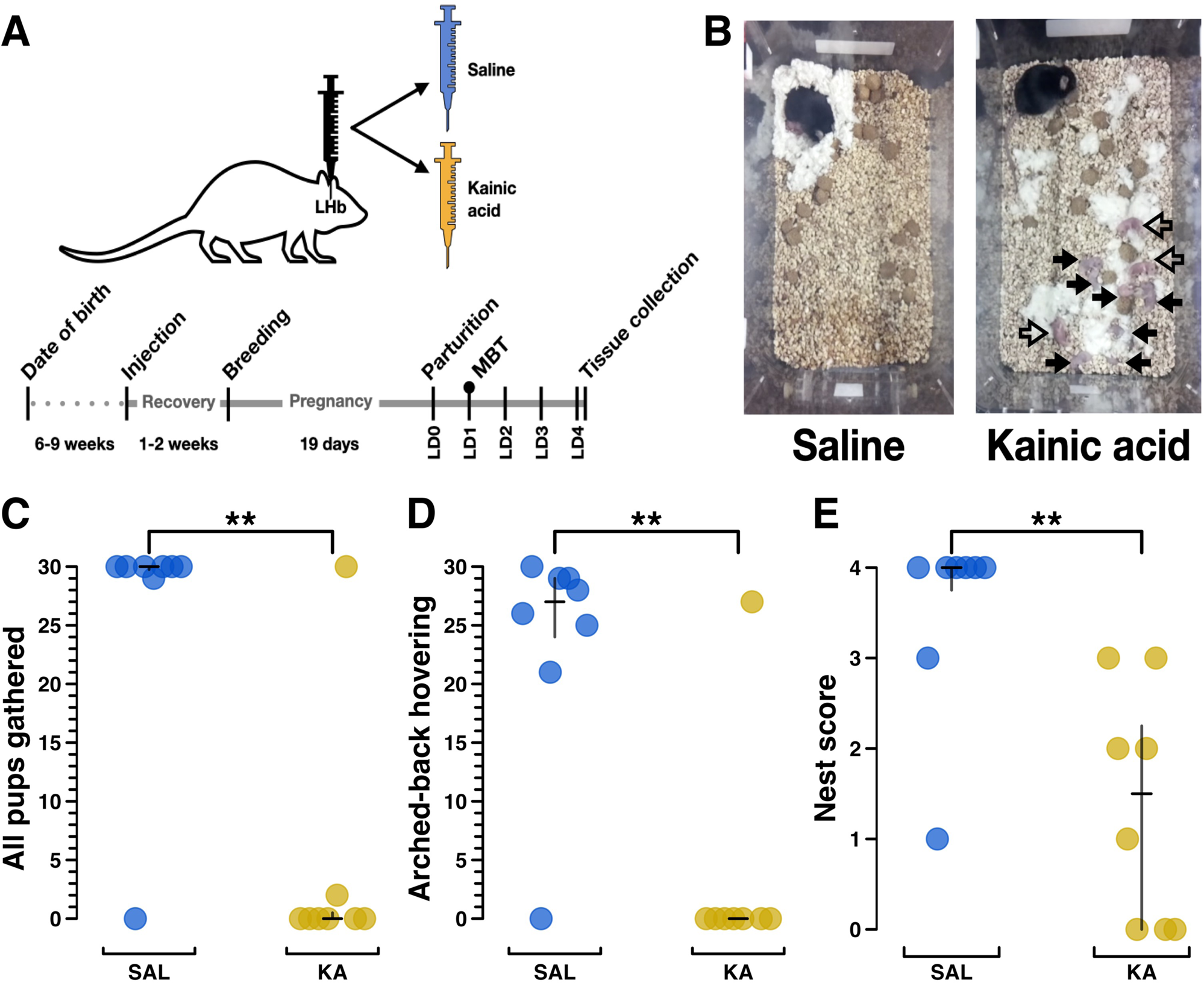
Kainic acid lesions of the lateral habenula (LHb) induce a severe maternal neglect phenotype in mice. A) Experimental schematic of surgery/injection treatment (top) and timeline (bottom). Female mice were injected bilaterally with either saline or kainic acid into the LHb. These mice were bred following recovery, and after pregnancy and parturition, behavioral data were collected on lactation day 1 (LD1), denoted by a circle above LD1 on the timeline. The experimental endpoint occurred on LD4, when dams were killed so that their brains could be collected for histology. B) Example still images showing the overhead view of a test cage containing a saline-treated and a kainic acid-treated female during a maternal behavior test (MBT). In the saline example, all pups are nested under the dam. In the kainic acid example, all pups are scattered. Filled arrows indicate dead pups and hollow arrows indicate living pups. C) Plot from MBT showing the total number of scored video windows (out of 30) in which each dam, indicated as a separate plot point, was observed with all pups gathered, for saline treatment (n=8) and kainic acid treatment (n=8). Note that one kainic acid treated mouse had no impairment in this score. Later histological analysis revealed that, for unknown reasons, this particular kainic acid injection failed to reduce the number or density of neurons in the LHb (see Figure 2). D) Plot from MBT showing total number of scored video windows (out of 30) in which each dam was observed engaged in arched-back hovering over any pups (presumed nursing) or was visibly nursing. E) Plot from MBT showing nest score (0: no nest attempted, 4: excellent nest, (Numan and Callahan, 1980)). Group differences were assessed using the Wilcoxon rank sum test, with significance levels of p < .05, p <.01, and p < .001 represented by *, **, and ***, respectively, here and for all subsequent figures. Median and interquartile range (IQR) are represented by black bars, here and for all subsequent figures. See Extended Figure 1-1 for further behavioral data.

### Lesion validation

Immunohistochemistry for NeuN enabled exhaustive manual counting of LHb neurons. This showed that saline-treated dams had about twice the LHb neurons of those dams treated with kainic acid (saline (n=6) median (IQR) = 9,239 (8615.75-9998), kainic acid (n=5) median (IQR) = 4171 (4111, 5149), *p* > .05) (Figure 2A, example images in Figure 2E). Neuronal density showed a similar pattern, with saline-treated dams having 2x the neuronal density of kainic-acid treated dams (saline (n=6) median (IQR) = 11.73 (10.51, 13.00), kainic acid (n=5) median (IQR) = 7.17 (6.43, 8.15), *p* > .05) (Figure 2B). While a reduction in LHb volume would not be surprising given the neuronal death associated with kainic acid lesion, any such difference was not statistically significant (saline (n=6), median (IQR) = 865 (725.75, 874.5), kainic acid (n=5), median (IQR) = 673 (582, 772), *p* = .0822) (Figure 2C).

**Figure 2.**
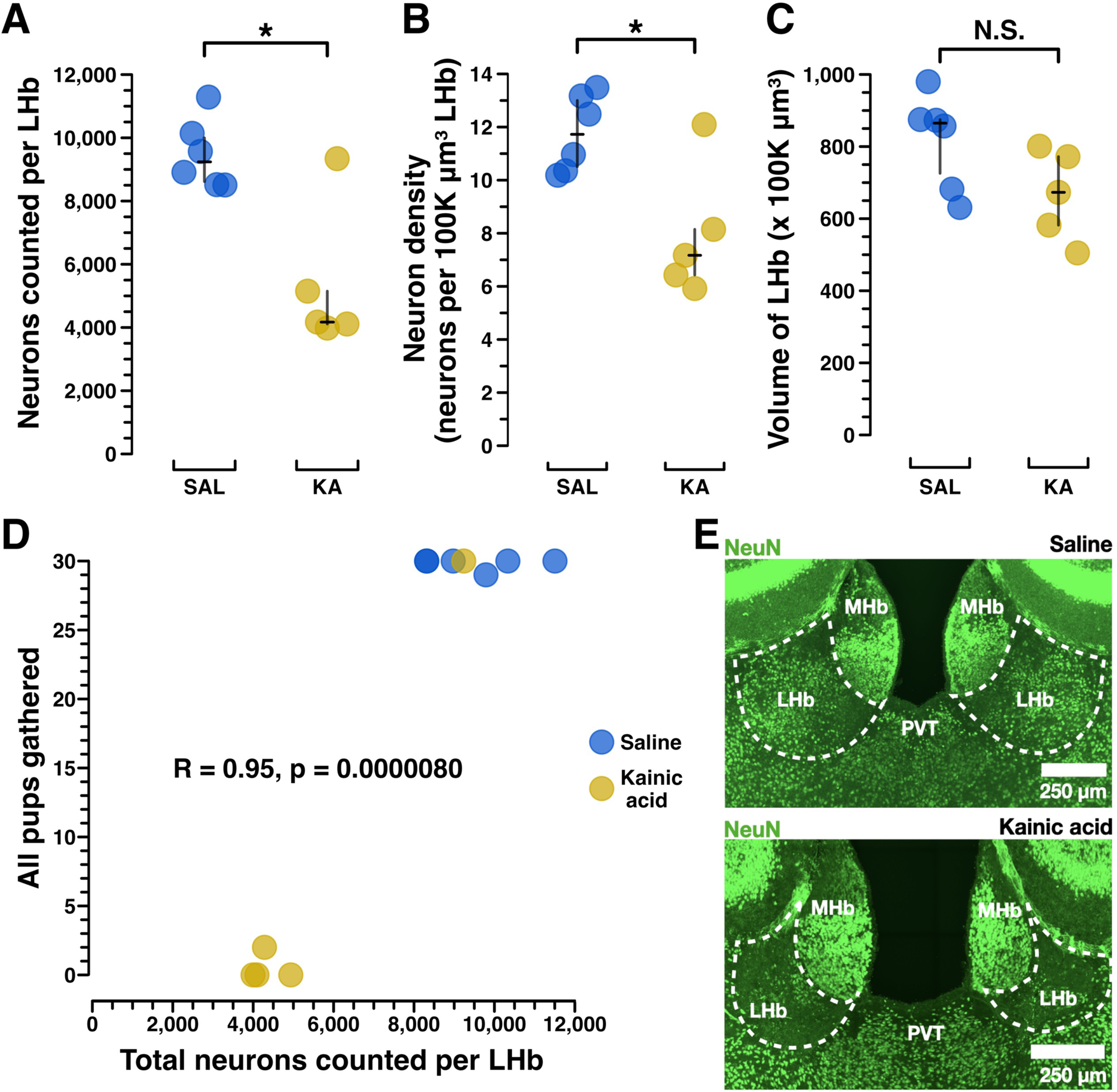
Kainic acid lesions reduced the number of LHb NeuN+ neurons. A) Manual NeuN+ LHb neuronal counting was performed to validate the lesions caused by kainic acid treatment versus saline treatment. Manual counting of NeuN+ neurons across the entire extent of LHb shows a significant reduction of NeuN+ neurons in all but one mouse. In mouse X306 kainic acid treatment failed to reduce the number of NeuN+ LHb neurons. Importantly, mouse X306 also mothered her pups indistinguishably from the saline treated group. Saline (n=6) median (IQR) = 9,239 (8615.75-9998), kainic acid (n=5) median (IQR) = 4171 (4111, 5149), *p* = .0303. B) To assess neuronal density, neurons per 100K µm^3^ LHb, a standardized unit of volume, was calculated. These results show kainic acid lesion reduced the NeuN+ neuronal density in LHb. Saline (n=6) median (IQR) = 11.73 (10.51, 13.00), kainic acid (n=5) median (IQR) = 7.17 (6.43, 8.15), *p* = .0303. C) Volumetric analysis of NeuN+ stained coronal sections across the anterior-posterior axis of LHb was conducted to see if total LHb volumes differed between groups, which could have occurred due to cell death following the lesion. Saline (n=6), median (IQR) = 865 (725.75, 874.5), kainic acid (n=5), median (IQR) = 673 (582, 772), *p* = .0822. D) Pearson correlation between MBT score for the “all pups gathered” measure, and total neurons counted per LHb shows that NeuN+ LHb neuron count is strongly correlated to behavioral performance of pup gathering. R = 0.95, *p* > .001. E) Exemplar confocal images of coronal sections (approximately −1.50mm posterior to bregma) show NeuN+ LHb neurons in a saline-treated dam and a kainic acid-treated dam. Statistical comparisons were performed using the Wilcoxon rank sum test.

Notably, the only kainic acid-treated dam that performed maternal behavior at control levels (mouse X306) was later shown to have the same number of total surviving neurons as those from the saline-treated group (Figure 2A, B, Extended Figure 7-2). The only saline-treated dam that neglected her pups (Figure 1C-E) may have reflected the known baseline rate of neglect in mouse dams (Schuster et al., 2023), as her brain could not be assessed for possible accidental injection damage, since the brain tissue from 5 animals (2 saline-treated and 3 kainic acid-treated), including hers, was damaged during COVID-19 lab shut downs and so the subsequent histology was uninterpretable.

Using behavioral results from only the brains with complete neuronal count data, we performed a Pearson correlation of dams’ “all pups gathered” scores and their total neurons counted per LHb (Figure 2D). This showed a correlation coefficient of *r* = .95 (*p* < 0.0001), suggesting a strong positive correlation between a dam’s surviving LHb neuron count and her maternal behavior performance

### Chronic chemogenetic LHb inactivation using DREADDs

The results from the kainic acid lesion experiments left open the possibility that LHb lesions blocked normal neuroplastic changes during pregnancy that are required for maternal behavior onset, but that the LHb may not be necessary for ongoing maternal behavior in the primiparous mouse dam. We sought to temporally refine the LHb manipulation through the use of inhibitory DREADDs. DREADDs are often used for cross-over experimental designs, wherein all animals are injected with a DREADD-expressing virus, but injection treatment is alternated each day within animals between saline and a DREADD-agonist. But maternal behavior is ill-suited to alternating saline days with DREADD-agonist days, since there are multiple latent variables, namely hormonally-mediated maternal behavior onset, learned maternal behavior acquisition, and changing pup needs, operating on a multiday timescale in the early postpartum period. Thus, we opted for chronic LHb inactivation (Benoit et al., 2022; Canetta et al., 2022; Tiwari et al., 2022) and injected the LHb of half the animals with an active inhibitory-DREADD virus (AAV2-hSyn-hM4D(Gi)-mCherry and the other half with a control virus (AAV2-hSyn-mCherry). From E17-LD4, all mice received 4x daily injections with Agonist-21, a designer ligand for the hM4D(Gi) metabotropic receptor (Figure 3A). Agonist-21 does not back-convert into psychoactive clozapine and has been shown to induce statistically significant behavioral effects as a result of DREADD agonism for 6 hours (Thompson et al., 2018; Jendryka et al., 2019; Ferrari et al., 2022). By administering Agonist-21 to both groups, the control group helps to control for the possibility of unknown off-target effects of Agonist-21.

**Figure 3.**
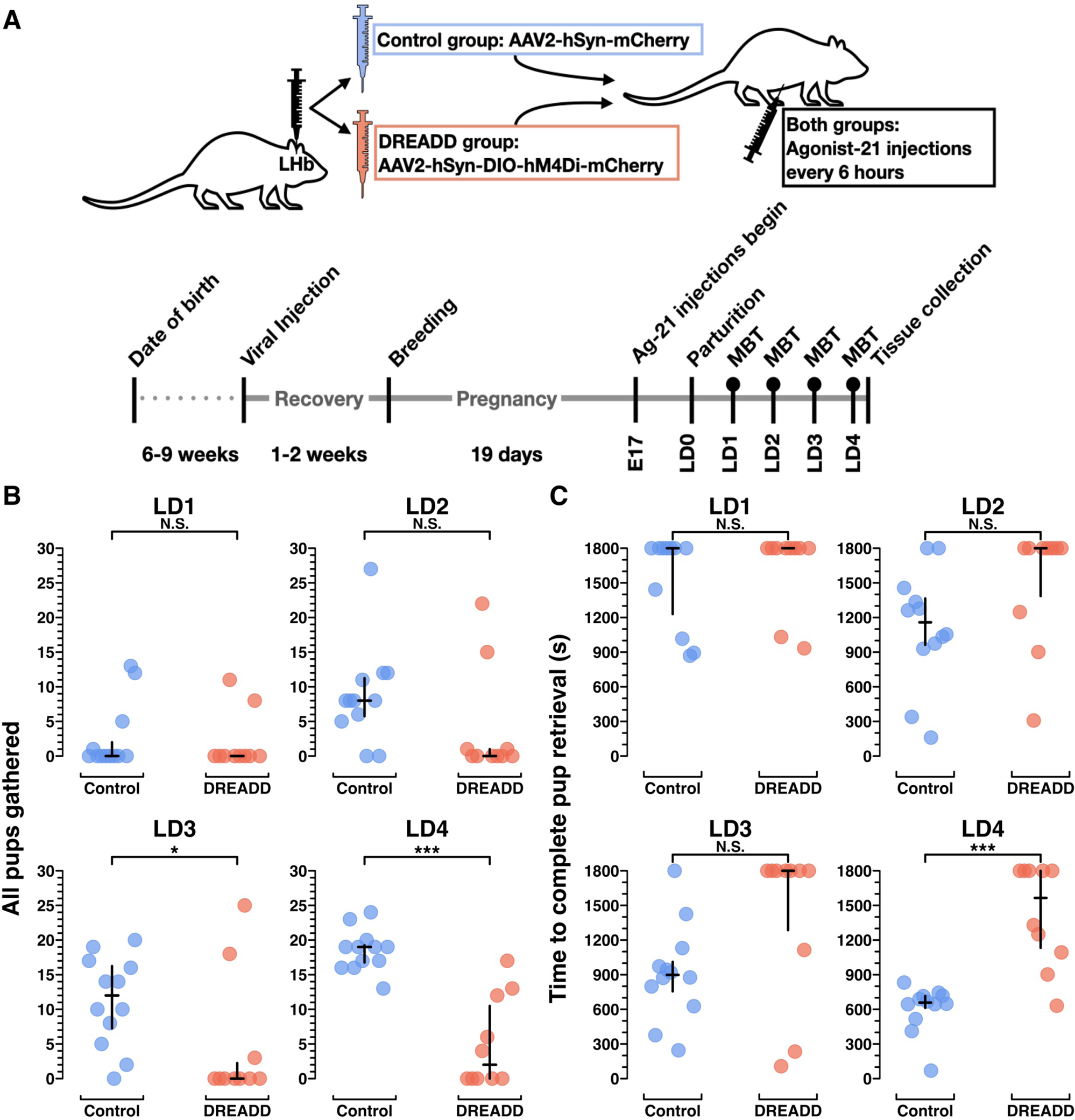
Chronic chemogenetic inactivation using DREADDs impairs the acquisition of pup retrieval, a novel maternal behavior, in primiparous dams. A) Experimental schematic of treatment (top) and timeline (bottom). Filled circles above timeline points Lactation Days 1-4 (LD1-LD4) denote behavior data capture for all chronic chemogenetic inactivation data presented elsewhere. B) Plot from MBTs LD1-LD4 showing total number of scored video windows (out of 30) in which each dam was observed with all pups gathered, after beginning the assay with all pups scattered. C) Plot from MBTs showing the number of seconds elapsed (of an 1800 sec long assay) to the point when the dam had completed retrieval of all pups. Statistical comparisons were performed using the Wilcoxon rank sum test. See Extended Figure 3-1 for further behavioral data.

### Chronic LHb inactivation impairs the acquisition of pup retrieval, a novel maternal behavior in primiparous dams

Since all dams in these experiments were pup-naïve until parturition, retrieving manually scattered pups to the nest was a novel behavior. Both control group dams and DREADD group dams were slow to retrieve their manually scattered pups on LD1 (*p* = .53), but by LD2 control group dams were already improving, while DREADD group dams performed similarly to the day before. By LD4, control group dams had learned to retrieve all of their pups at least 2-3x faster than DREADD groups dams (*p* > .001) (Figure 3B, C; Extended Figure 3-1). On LD4, DREADD dams were divided bimodally, with five DREADD dams still failing to complete pup retrieval at all during the 30-minute assay (and thus receiving a score of 1,800), and the other five DREADD dams eventually completing pup retrieval, yet doing so much slower than the control group dams (*p* > .001) (Figure 3C, Extended Figure 3-1). For example videos of LD4 behavior assays, see Extended Figure 3-2, and 3-3. Altogether, these data suggest chronic LHb inactivation blocked acquisition of the novel pup retrieval behavior completely in about half the DREADD group, and at minimum impaired acquisition in all but 2 of the DREADD females (n=10) by LD4 (Figure 3B,C, Figure 5A).

### Chronic LHb inactivation reduces nest building, an established behavior in primiparous dams

Unlike rats, virgin adult mice spontaneously build nests (Oortmerssen, 1971; Van Loo and Baumans, 2004). As such, primiparous mouse dams do not need to learn to build nests. But since highly-motivated nest building and high nest scores in pregnant female mice predict engagement in other maternal behaviors as well as pup survival (Schuster et al., 2023), nest building motivation is an important maternal behavior variable.

Across all days, control group dams spent significantly more time engaged in nest building than DREADD group dams (Figure 4A, Extended Figure 3-1). Control group dams also built nests more reliably than DREADD group dams, with higher overall nest scores (Figure 4B, Extended Figure 3-1). But, similar to pup retrieval, DREADD dams’ nest scores on LD4 were bimodal, with about half the DREADD group females receiving high nest scores, and the other half failing to build a nest at all. These results suggest that for about half the DREADD group, LHb inactivation severely impaired nest building motivation.

**Figure 4.**
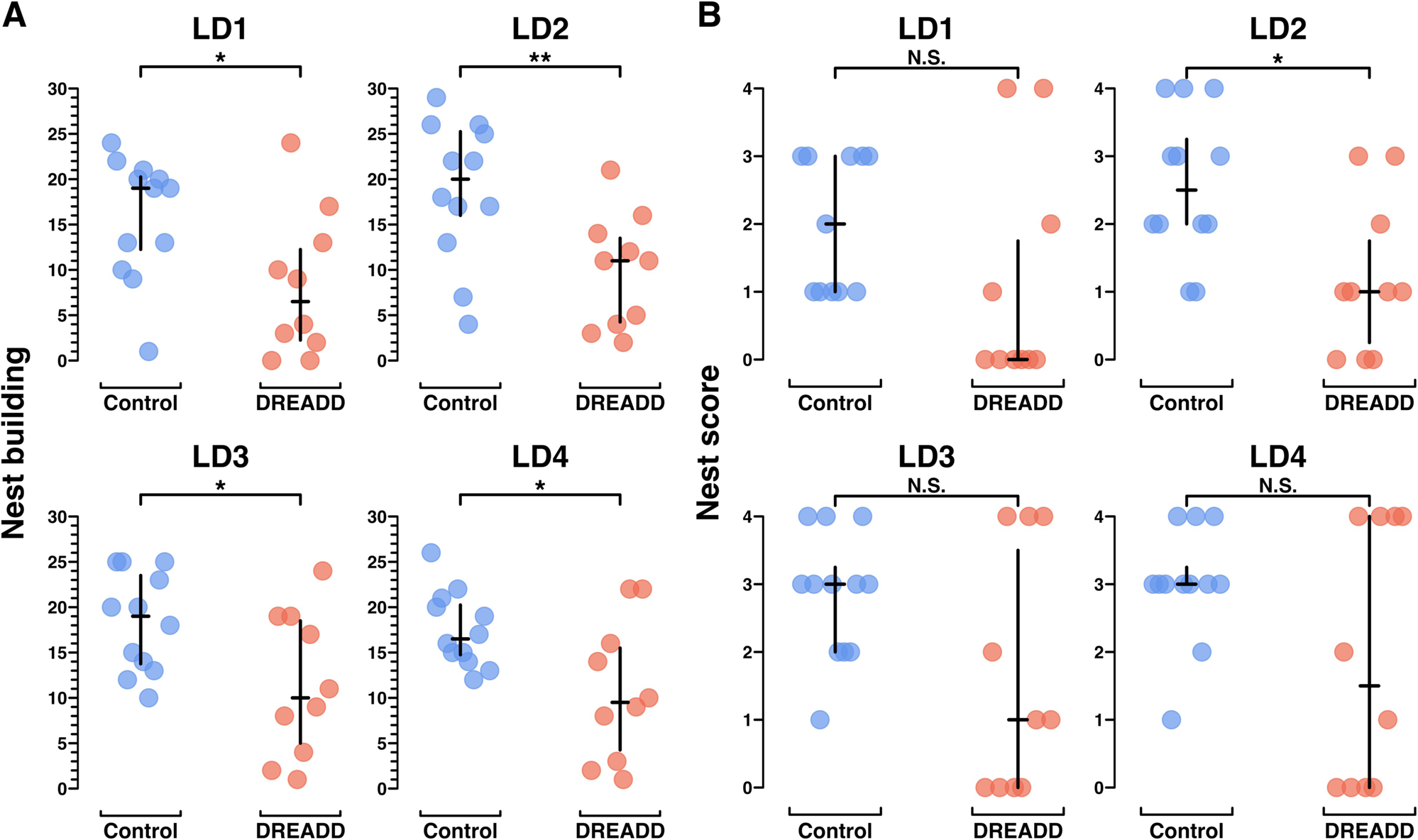
Chronic chemogenetic inactivation decreases nest building behavior in primiparous dams. A) Plots from MBTs LD1-LD4 showing the total number of scored video windows (out of 30) each dam was observed engaged in nest building. Unlike pup retrieval, nest building is an established behavior in virgin female mice. B) Plots from MBTs LD1-LD4 showing the nest score at 30 minutes. Statistical comparisons were performed using the Wilcoxon rank sum test. See Extended Figure 3-1 for further behavioral data.

### Linear mixed model of a novel versus an established maternal behavior in primiparous mouse dams

Since each univariate analysis presented in Figure 3 and 4 is analyzed in isolation of any other lactation day, significance testing only assesses group differences at that individual time point. Since what we could gather visually from Figure 3, that there appeared to be impaired learning of pup retrieval in DREADD-group females, was not being tested by the Wilcoxon rank sum tests, we sought a superior statistical approach. Specifically, we sought an approach that would enable analysis of each dam’s individual longitudinal performance, nested within her treatment group identity. In consultation with the [redacted for double-blind review] Biostatistics Center, we conducted a random intercept analysis, a type of random effects mixed linear model (Verbeke, 1997; Barr, 2013; Bates et al., 2015). Random intercept models enable analysis of multivariate interactions between treatment group and behavior variables, across time or space, which enables each datapoint to be linked to the individual animal to which it belongs. The model has the fixed effects, which are the effects of the parameters of the experiment (i.e. “All pups gathered”, lactation day, treatment group, and animal ID for Figure 5B). As an additional improvement over univariate methods, the model also has random effects (i.e. individual animal IDs) introduced by random individual differences between mice or their particular injection surgeries. This enables contextualization of the effect size of the tested interaction, in comparison to any random effects introduced by non-independent terms. This form of analysis is widely used in adjacent fields, and based on conversations with several biostatisticians, ought to be being used every time, for example, results from multiple cohorts of animals are analyzed together. Put another way, anytime there is hierarchical structure to data, like if thousands of cells are sequenced across 12 animals, those data need to be pooled in a manner accounting for the non-independence of the data from each animal.

**Figure 5.**
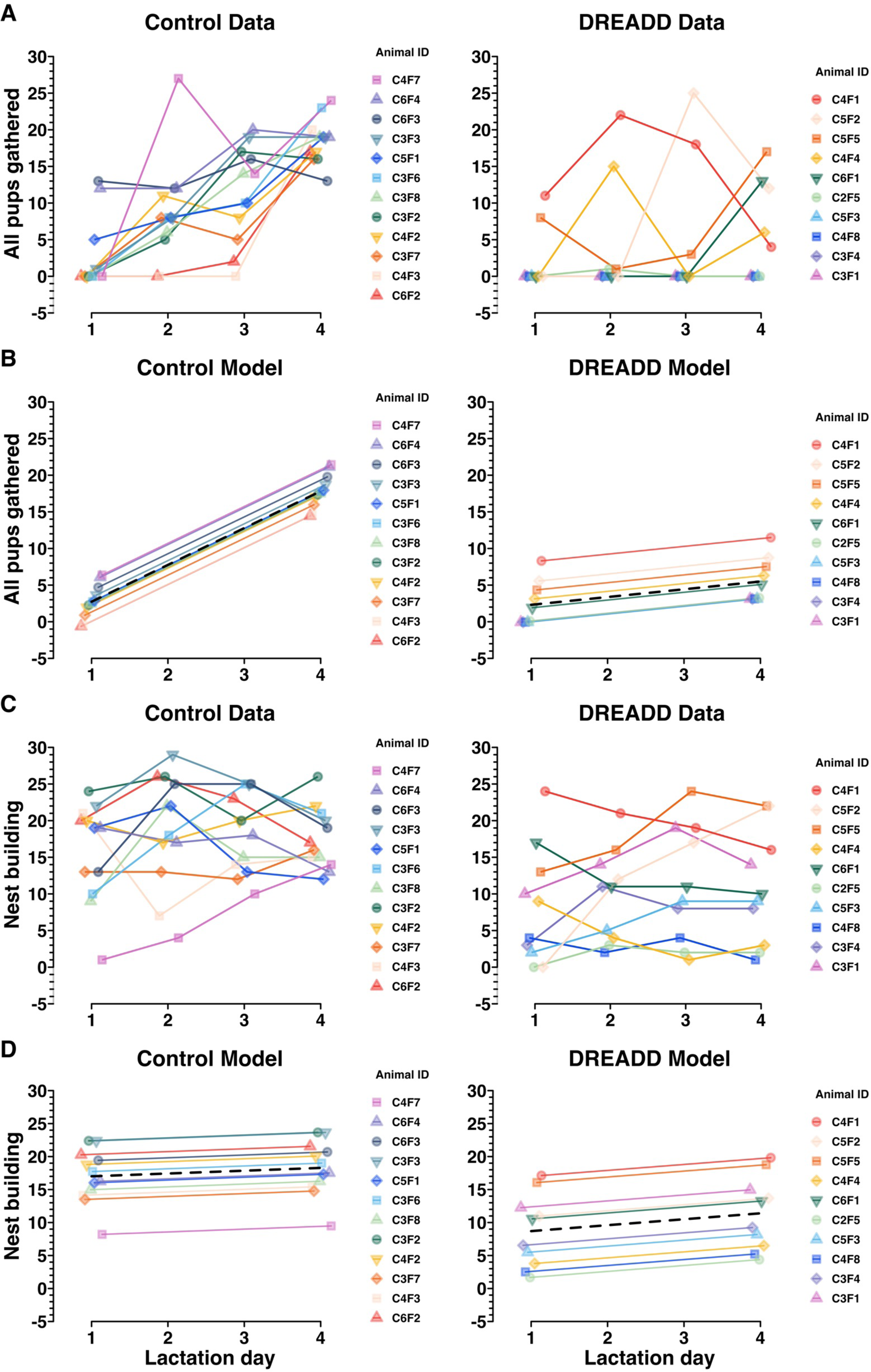
The effect of chronic chemogenetic inactivation on retrieval and nesting behaviors was compared across time and between groups, assessed using a random intercepts linear mixed model. A) Control data and DREADD data (also depicted in Figure 3B) showing the effect of time on the “all pups gathered” variable in MBTs. A positive slope suggests that an animal acquired the novel maternal behavior of pup retrieval to gather all of her pups over the timescale of the experiments. B) A random intercept linear mixed model depicting control group and DREADD group trajectories (black dashed lines), and individual dam trajectories across time for the “all pups gathered” variable. This model assesses whether the passage of time affects the groups differently on the “all pups gathered” variable (indicated by a difference between group slopes) and estimates the random individual differences for each animal. C) Control data and DREADD data (also depicted in Figure 4A) showing nest building MBT performance over the timescale of the experiments. D) A random intercept linear mixed model depicting control group and DREADD group trajectories (black dashed lines) and individual dam trajectories for each group across time for the “nest building” variable. As expected, no effect of time on nest building behavior in MBTs was found. Since nest building is an established behavior in virgin female mice, the group intercept is of greater interest to assess a group difference in performance on the nest building variable. See Extended Figure 5-1 for exact term values, confidence intervals, degrees of freedom, *t* values, and significance testing.

In Figure 5A, the raw behavior data for “all pups gathered” shows each dams’ performance for each MBT (LD1-LD4), with one plot for each treatment group. That same data is then modeled in Figure 5B, where the random intercept model finds a statistically significant (*p* > .001) difference between the slopes for each group (Extended Figure 5-1). No significant difference is found between group intercepts (*p* = .26), suggesting there is no initial difference in retrieval motivation or performance between control and DREADD groups, but only the difference that emerges over time, as the control group dams learn to conduct pup retrieval while the DREADD group dams’ pup retrieval learning is impaired (Figure 5, Extended Figure 5-1).

In Figure 5C and D the raw data and modeled data are shown for nest building performance over time. In contrast to the random intercept model of retrieval, nesting behavior is already established and thus both groups have very subtle slopes with no between group difference (*p* = .57). There is, however, a statistically significant difference in group intercepts, with the control group engaging in twice as much nest building as the DREADD group, across time (*p* > 0.01) (Figure 5D, Extended Figure 5-1). This difference suggests an intergroup difference in baseline nest building motivation.

### mCherry immunoreactivity in dams’ brains following chronic chemogenetic inactivation behavior experiments

We sought to confirm viral injection targeting and to assess whether maternal behavior was differentially impacted by the number of mCherry+ LHb cells expressing DREADDs. mCherry+ LHb cells were exhaustively counted across both groups and total LHb volumes were measured (Figure 6A, C). The control group virus had slightly better transfection efficiency than the DREADD group virus (Figure 6B). Counter to the results seen in the kainic acid lesion validation data (Figure 2), the number of mCherry+ LHb cells did not predict LD4 performance in DREADD group dams for either pup retrieval (Extended Figure 6-1A) or nest building (Extended Figure 6-1B). In fact, many of the dams with severely impaired pup retrieval had quite modest LHb transfections. This finding raises the questions of whether a neighboring brain region with off-target DREADD expression could be predictive of maternal behavior, and whether particular LHb subregions or cell types may be most relevant for maternal behavior regulation.

**Figure 6.**
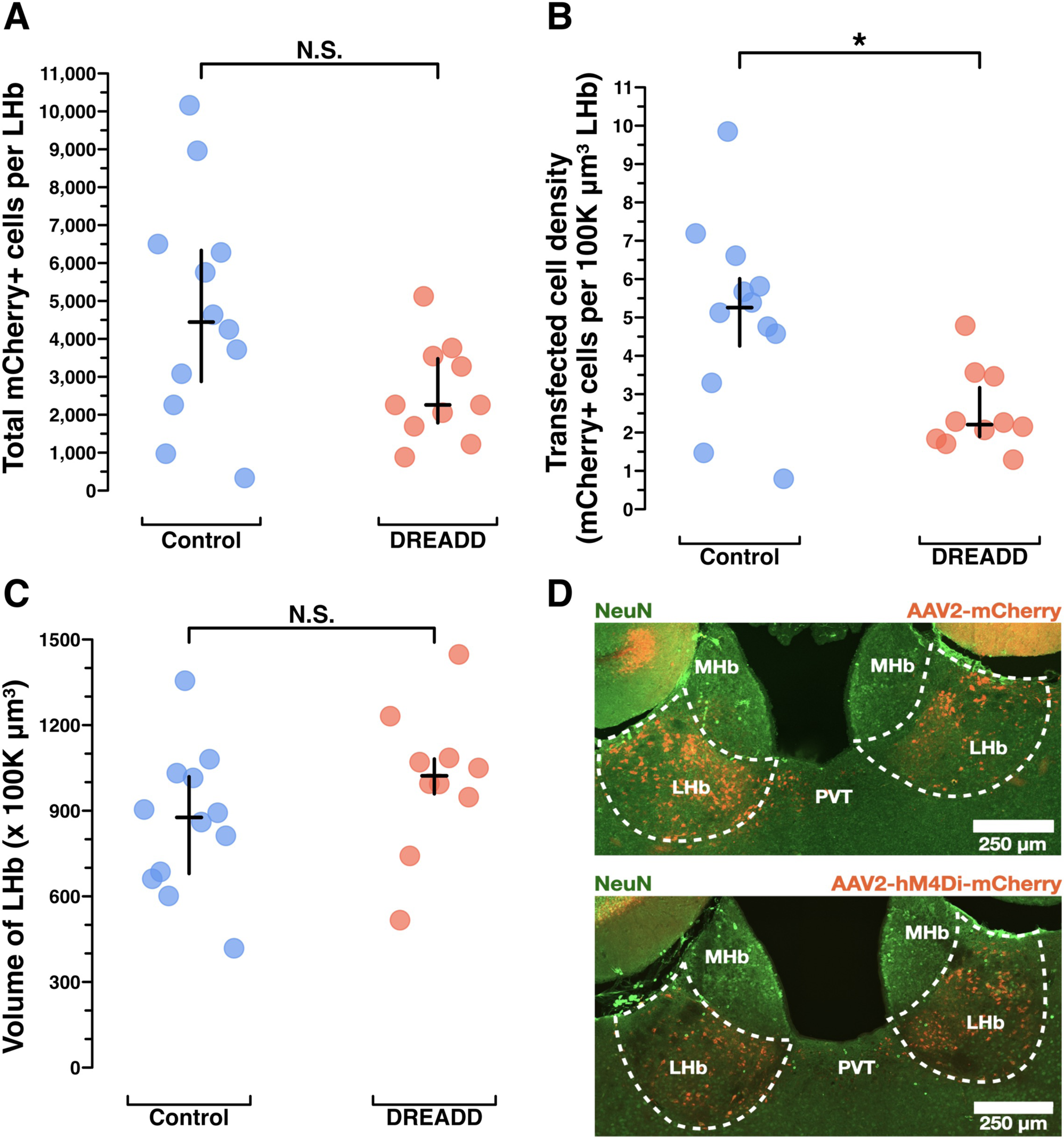
Histological analysis of mCherry immunoreactivity to assess DREADD transfection efficiency in dams’ brains following chronic chemogenetic inactivation behavior experiments. A) Results from manual counting of mCherry+ cells across the entire extent of LHb show control dams likely had more mCherry+ LHb cells than DREADD-treated dams, see results section for more detail. Controls (n=12) median (IQR) = 4,444 (2876, 6334), DREADDs (n=10) median (IQR) = 2258 (1785, 3478), *p* = .0648. B) Results from manual volumetric analysis of coronal sections across the entire anterior-posterior axis of LHb to allow for density calculations in C. Controls (n=12) median (IQR) = 876.5 (679.70, 1019.10), DREADDs (n=10) median (IQR) = 1022.30 (959.10, 1081.30), *p* = .1402. C) Results for mCherry+ cells per 100k µm^3^ LHb as a standardized unit of volume show control dams had a higher density of mCherry+ LHb cells than DREADD-treated dams. Controls (n=12) median (IQR) = 5.257 (4.256, 6.011), DREADDs (n=10), median (IQR) = 2.205 (1.893, 3.169), *p* = .0169. D) Example histology of mCherry+ LHb cells in a control dam and a DREADD dam (approximately −1.50mm posterior to bregma). Statistical comparisons were performed using the Wilcoxon rank sum test.

To address the question of whether off-target transfections predict maternal behavior in DREADD group dams, a blinded scorer assigned subjective scores to each brain section of hippocampus, medial habenula, paraventricular nucleus of the thalamus, central lateral nucleus of the thalamus, and anterior pretectal nucleus. Spearman correlations showed none of the off-target regions’ transfection magnitudes was related to DREADD dams’ performance on LD4 for “all pups gathered” or “nest building” (Extended Figure 6-2).

### Spatial histology random intercepts model

To partially answer the question of whether particular LHb subregions may be most relevant for maternal behavior, we performed another random intercept model, this time of the kainic acid lesion histology data (Figure 7). Since we sought to test whether the spatial distribution of surviving neurons along the anterior-posterior (A-P) axis differed by maternal behavior outcome, we excluded dam X306 (the single kainic acid-treated dam who mothered her pups, Figure 1C-E, Figure 2D) from this analysis so that within-group maternal behavior outcomes were homogenous. This was done in concordance with the recommendations from [redacted for double-blind review].

**Figure 7.**
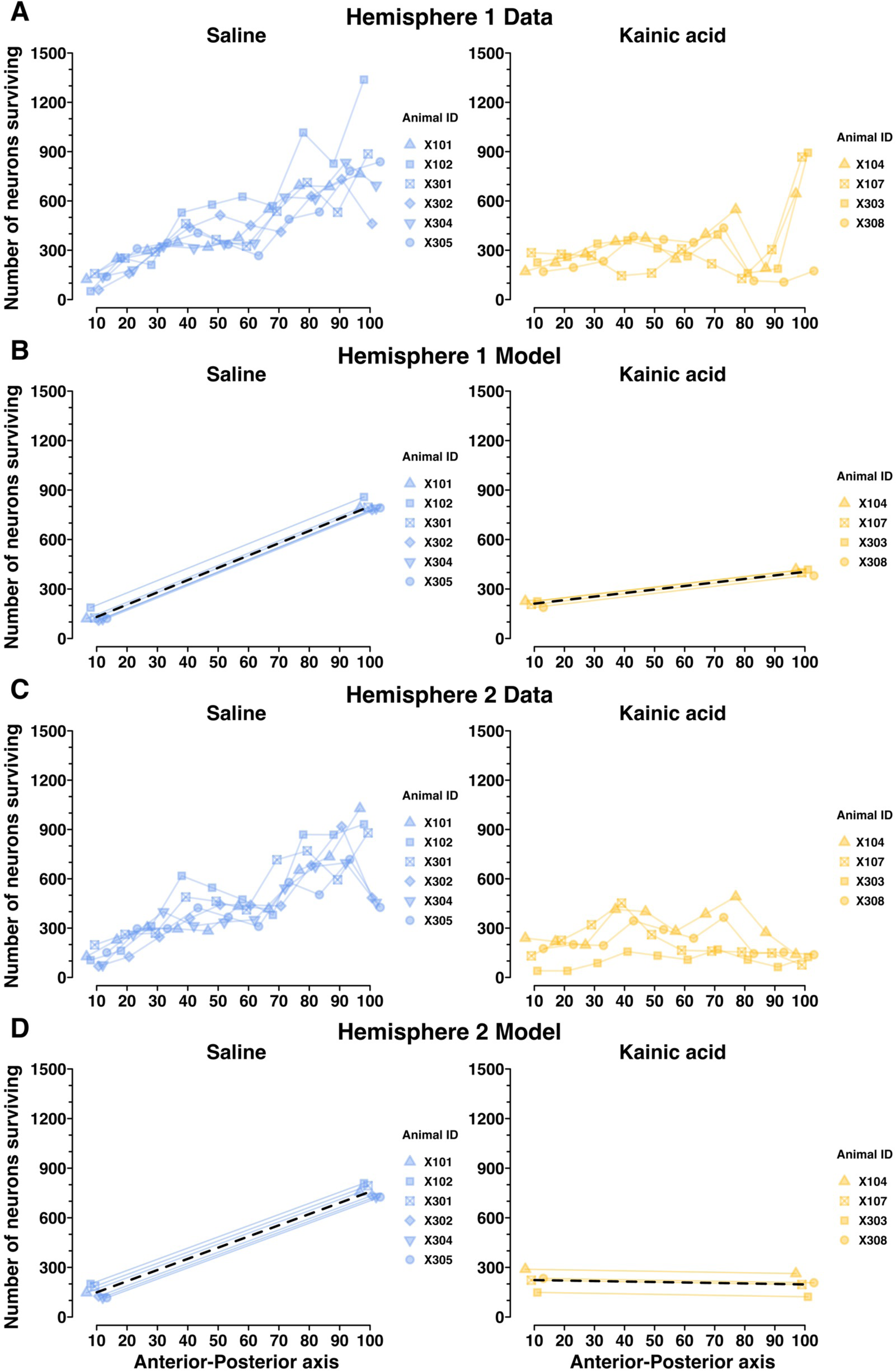
The LHb neurons regulating maternal behavior may preferentially reside in the posterior LHb, assessed using a random intercepts linear mixed model. Here, we test whether the spatial distribution of surviving neurons along the AP axis of LHb is significantly related to maternal behavior outcomes. A) NeuN+ LHb neuron counts in Hemisphere 1 for each decile of the AP axis of LHb. B) A random intercepts linear mixed model depicting saline-treated and kainic acid-treated group trajectories, and individual dam trajectories, for NeuN+ LHb neuron counts in Hemisphere 1 across the AP axis. This model, along with that in D, shows a statistically significant relationship between maternal behavior outcomes and LHb neuron counts by AP axis decile, and suggests that the maternally-relevant LHb neurons may reside at the posterior aspect of LHb. C) NeuN+ LHb neuron counts in Hemisphere 2 for each decile of the AP axis of LHb. D) A random intercepts linear mixed model depicting saline-treated and kainic acid-treated group trajectories, and individual dam trajectories, for NeuN+ LHb neuron counts in Hemisphere 2 across the AP axis. This again shows a statistically significant relationship between maternal behavior outcomes and LHb neuron counts by AP axis decile, particularly suggesting the posterior aspect of LHb may be the most important for maternal behavior. See Extended Figure 7-1 for model terms, confidence intervals, degrees of freedom, t values, and significance testing.

With the remaining 10 dams’ histology data (n = 4 kainic acid; n = 6 saline), we segmented surviving neuron counts by decile along the A-P axis. The 10^th^ percentile marks the most anterior tenth of LHb sections quantified, and the 100^th^ percentile marks the most posterior tenth of LHb sections quantified (Figure 7, Extended Figure 7-1). Each hemisphere was analyzed independently. The spatial histology data for Hemisphere 1 and Hemisphere 2 are presented in Figure 7A and 7C, respectively. Those same data are then modeled in Figure 7B and 7D. The random intercept model finds no statistically significant difference between groups at the 10^th^ percentile AP mark (*p* = .17 and *p* = .19 for Hemispheres 1 and 2), but finds a significant difference between groups at the 100^th^ percentile A-P mark, and between group slopes from 10^th^ to 100^th^ percentiles, for both hemispheres (*p* > .001 for 100^th^ percentile mark, and group slopes, for both hemispheres) (Extended Figure 7-1). In the table presented in Extended Figure 7-1, significance testing for saline-treated group-term values (i.e. intercepts at 10^th^ and 100^th^ percentile AP mark and group slope) reflect the model’s confidence that the true saline-group term-value is nonzero, whereas the significance testing for the kainic acid-treated group terms reflect the model’s confidence that there is a significant difference between saline and kainic acid groups. This is all to say, the results of the significance testing in Extended Figure 7-1 that tests for differences between the two groups, are the p-values printed in the rows for labeled “Kainic acid” group. These findings suggest that surviving neuron counts in the posterior LHb, and not the anterior LHb, are predictive of maternal behavior outcomes.

Animal X306 was kainic acid-treated but went on to mother her pups indistinguishably from dams in the saline-treated group. Her total surviving LHb neuron count was also indistinguishable from those of the saline group (Figure 2A). However, upon reviewing her spatial histology, X306 has lower neuron counts in the anterior-most deciles of LHb compared to saline dams. This could mean her kainic acid injection failed, and her anterior LHb neuron counts are the result of normal variation. Or, her lesion had an anterior LHb bias, thus sparing the central and posterior LHb. She displayed intact maternal behavior. Although no conclusions can be drawn from her single case, X306’s spatial histology and maternal behavior outcome support our principal spatial histology finding that an intact posterior LHb enables maternal behavior. We present her individual spatial histology data separately for comparison (Extended Figure 7-2).

## Discussion

The goal of the present study has been to determine whether the LHb is required for maternal behavior in the naturally parturient primiparous mouse dam. We found that lesioning the LHb with kainic acid induced a severe maternal neglect phenotype in the mouse dam. Lesioned dams ignored their pups completely, failing to ever feed or retrieve them to the nest. Because of this neglect the pups all died within 60 hours of birth (Figure 1, Extended Figures 1-1, 1-2, 1-3).

To test the hypothesis that ongoing LHb activity is required for maternal behavior, we chronically inactivated LHb using DREADDs starting just before parturition. This experiment helped exclude the possibility that kainic acid lesions caused the maternal neglect phenotype by blocking neuroplastic changes during pregnancy from occurring, rather than because LHb function is actually necessary to ongoing maternal behavior in the parturient mouse dam. To compare the effect of LHb inactivation on novel maternal behaviors and established behaviors in primiparous dams, pup retrieval (after scattering by the experimenter) and nest building were examined (Figures 3 and 4, Extended Figure 3-1, 3-2, 3-3) and analyzed with a random intercepts linear mixed model (Figures 5, Extended Figure 5-1). DREADD-treated dams failed to learn to retrieve their pups across MBTs performed on LD1-LD4. The control-treated dams learned to retrieve their own pups over the same period. Simultaneously, nesting behavior was suppressed in DREADD-treated dams compared to control dams.

Finally, we conducted a spatial analysis comparing surviving NeuN+ LHb neuron counts in kainic acid-treated and saline-treated dams along the anterior-posterior axis. We found that surviving NeuN+ neuron counts in the posterior element of LHb were most predictive of maternal behavior outcomes (Figure 7, Extended Figures 7-1 and 7-2). The animals included in our analysis had central- and posterior-biased LHb lesions (Figure 7, Extended Figure 7-1), so this leaves open the possibility a large sample of anterior-restricted LHb lesions could produce similar results. Our sole data point suggesting anterior-restricted LHb lesions do not impact maternal behavior is inconclusive due to lack of sample size (Extended Figure 7-2). So, future experiments would benefit from specific targeting of anterior versus posterior LHb.

Random intercept linear mixed models were utilized because in both the chronic chemogenetic inactivation experiments and the kainic acid spatial histology data, the data were hierarchical in nature (each animal contributed multiple data points). The mixed linear models enable the multivariate analysis of hierarchical data (in this case, across time in Figure 5 and across space in Figure 7; for more see results section or Verbeke, 1997; Barr, 2013; Bates et al., 2015).

Here, we establish that the LHb is required for maternal behavior in the naturally parturient mouse dam. Work from the Morrell lab in the 1990s examined the role of the LHb in maternal behavior in the rat dam (Matthews-Felton et al., 1995) but the findings have yet to be extended to the mouse dam, nor the mouse dam studied in conjunction with behavior towards her own biological pups. Virgin female mice are not innately pup-avoidant like virgin female rats and through sensitization exhibit a similar array of parental behaviors to the parturient mouse dam (Leblond, 1938; Kohl and Dulac, 2018). This leads sensitized virgin female mice to frequently be used, instead of lactating dams, to study mouse parenting behavior. But virgin mouse females’ parental behavior acquisition, hormonal environment, gene expression, and motivational state all differ significantly from those of naturally parturient dams (Gandelman et al., 1970; Hauser and Gandelman, 1985; Abellán-Álvaro et al., 2021; Carcea et al., 2021). By establishing the role for the LHb in maternal behavior in the naturally parturient mouse dam, we solidify the LHb as a component of shared parental circuitry between alloparental virgin females and naturally parturient mouse dams.

One alternative explanation of our findings would be an adjacent off-target area incidentally lesioned with kainic acid or transfected with DREADD virus being responsible for the maternal neglect phenotype. We find this possibility unlikely, since kainic acid lesioned animals’ neighboring brain regions were closely inspected for loss of NeuN+ neurons and none was found. In the chemogenetic inactivation experiments off-target transfections were fairly common. Off-target transfections were particularly prevalent in hippocampus, which was located along the injection trajectory for LHb, and in the anterior pretectal nucleus, which was highly sensitive to transfection by trace amounts of virus. We verified there was no relationship between any off-target transfection magnitudes and maternal behavior outcomes by conducting blinded subjective scoring of mCherry+ cells in off-target regions and analyzing the results in relation to retrieval and nesting behavior in DREADD-treated dams. None of the off-target transfections versus maternal behavior analyses approached significance (Extended Figure 6-2). Together with the spatial histology data from the kainic acid lesions (Figure 7, Extended Figure 7-1), we feel confident the effect of maternal behavior from these manipulations arises from perturbations of the LHb.

During DREADD surgeries, we had no notion as to a particular zone of interest along the AP axis of LHb, so we injected each animal with multiple viral injections along the AP axis in an attempt to ensure anterior, central, and posterior LHb were all transfected. Future work could use single bilateral viral injections targeted to posterior LHb to greatly limit off-target transfections. The experimental design of the chemogenetic inactivation experiments did not enable spatial histology analysis since dividing the DREADD-treated dams into maternal behavior-outcome-based groups was non-trivial given the heterogeneity of some dams’ maternal behavior (such as dams who engaged in retrieval but not nesting, or the reverse) (Figure 5).

The severe maternal neglect phenotype following kainic acid lesion and the maternal behavior deficits caused by chronic LHb inactivation are likely related to reward signaling in the regulation of maternal behavior. Maternal behavior, like all motivated behavior, requires tracking ‘value changes’, e.g. “a pup has just fallen out of the nest”, and ‘value states’, e.g. “that pup’s distress call is aversive,” to motivate flexible decision-making, e.g. a dam retrieving the displaced pup to the nest (Proulx et al., 2014). Through input from the ventral pallidum and reciprocal connectivity with the serotonin and dopamine systems, the LHb plays an important role in the integration of value changes and value states (Reisine et al., 1982; Amat et al., 2001; Matsumoto and Hikosaka, 2007; Yang et al., 2008; Kim, 2009; Omelchenko et al., 2009; Bromberg-Martin and Hikosaka, 2011; Stamatakis and Stuber, 2012; Jhou et al., 2013; Stamatakis et al., 2013; Amo et al., 2014; Brown et al., 2017; Stephenson-Jones et al., 2020; Lalive et al., 2022; Mondoloni et al., 2022).

There is strong evidence of spatial segregation of LHb cell types. Geisler et al. (2003) conducted immunohistochemical analysis of 30 antibody markers in rat habenular sections and found specific subnuclei marked by neurofilament H, Kir3.2 protein, GABA B receptor and tyrosine hydroxylase. Quina et al. (2015) performed extensive LHb-efferent tracing in the mouse, using Pou4f1 antibodies to label the posterior two thirds of LHb, and finding retrograde median raphe injections preferentially label the medial division of LHb. Two single-cell RNA-sequencing and *in situ* hybridization of the mouse habenula projects were undertaken by two separate labs (Hashikawa et al., 2020; Wallace et al., 2020) and showed substantial agreement, including large populations of cells with transcripts for Pcdh10, Gpr151, Chrm3 and Gap43, all of which are spatially divided. Gap43 shows topographic overlap with the neurons of the medial division that Quina et al. (2015) found to project to median raphe. VTA-dopamine neurons seem to preferentially receive input from Chrm3+ LHb cells (Wallace et al., 2020). Necab1 and Cartpt represent still other spatially organized cell populations in LHb (Hashikawa et al., 2020). Thus, future work could conduct in vivo recordings of LHb cells with specific genetic markers or projection targets, such as Gap43 or the median raphe, which seem to largely receive input from neurons along the medial division of LHb, and record neural activity during maternal behavior. This would allow analysis of differential involvement of a given neuronal subpopulation in the established nest building behavior versus novel maternal behaviors of pup retrieval and crouching behaviors. This could build on prior findings that nest building may be genetically independent from pup retrieval and crouching which may be regulated together by genes at the same loci (Bendesky et al., 2017). In the same animals, neural responses to perturbations of value changes and value states could be assessed, enabling significant progress towards understanding the transformation of pup-related stimuli into maternal behavior in the mouse dam.

Given our findings that the LHb is required for maternal behavior in the naturally parturient mouse dam, and the wealth of evidence that the LHb, as a critical regulator of reward signaling, is also involved in the pathophysiology of depression and anxiety (Matsumoto and Hikosaka, 2007; Sartorius and Henn, 2007; Yang et al., 2008, 2018; Li et al., 2011; Proulx et al., 2014; Shabel et al., 2014), we believe there is substantial evidence to warrant the investigation of the LHb as a possible target for future therapeutics in the treatment of postpartum depression and anxiety (Li, 2022).

## Acknowledgements

We’d like to acknowledge the contributions of Carol B. Thompson, of the Johns Hopkins Institute for Clinical and Translational Research. This publication was made possible by the Johns Hopkins Institute for Clinical and Translational Research (ICTR) which is funded in part by Grant Number UL1 TR003098 from the National Center for Advancing Translational Sciences (NCATS) a component of the National Institutes of Health (NIH), and NIH Roadmap for Medical Research. Its contents are solely the responsibility of the authors and do not necessarily represent the official view of the Johns Hopkins ICTR, NCATS or NIH. We’d also like to acknowledge the truly generous mentorship of Dr. Surjit Chandhoke, Dr. Manavi Chatterjee, Dr. Solange Brown, and Dr. Keri Martinowich.

