## Extended Figure 1-1 for "The lateral habenula is required for maternal behavior in the mouse dam"

| Variables (n of 30 scored video windows) | Kainic acid injected dams (n=8) | | Saline injected dams (n=8) | | *p* value | |
|---|---|---|---|---|---|---|
| | Median | (IQR) | Median | (IQR) | | |
| All pups gathered | 0 | (0, 0.5) | 30 | (29.75, 30) | 0.0097 | ** |
| Arched-back hovering/visible nursing | 0 | (0, 0) | 27 | (24, 29) | 0.007 | ** |
| Time with pups | 0 | (0, 0) | 10.5 | (5, 14.25) | 0.0098 | ** |
| Pup interactions ‡ | 0.5 | (0, 1) | 6 | (4.5, 12.5) | 0.0213 | * |
| Nest building | 0.5 | (0, 2.25) | 2 | (1, 3) | 0.3067 | |
| Solo activity | 11.5 | (4, 15.25) | 5 | (1, 6.25) | 0.0824 | |
| Solo rest | 16 | (11.25, 22.25) | 0 | (0, 0.25) | 0.0184 | * |
| Pup lacking milk spot visible | 30 | (29.75, 30) | 0 | (0, 0) | 0.0067 | ** |
| Dead pup visible | 14.5 | (0, 30) | 0 | (0, 0) | 0.0124 | * |
| Nest score (0-4) | 1.5 | (0, 2.25) | 4 | (3.75, 4) | 0.0047 | ** |
