## Extended Figure 3-1 for "The lateral habenula is required for maternal behavior in the mouse dam"

| Variables | Lactation Day (LD0 is DOB) | Control dams AAV2-hSyn-mCherry (n=12) Median | (IQR) | DREADD dams AAV2-hSyn-hM4Di-mCherry (n=10) Median | (IQR) | *p* value | |
|---|---|---|---|---|---|---|---|
| All pups gathered | | | | | | | |
| (n of 30 scored video windows) | LD1 | 0.00 | (0.00, 0.00) | 0.00 | (0.00, 0.00) | .5286 | |
| | LD2 | 8.00 | (5.75, 11.25) | 0.00 | (0.00,1.00) | | |
| | LD3 | 12.00 | (7.25, 16.25) | 0.00 | (0.00,2.25) | | |
| | LD4 | 19.00 | (16.75, 19.25) | 2.00 | (0.00,10.5) | .0000 | *** |
| All pups nested | | | | | | | |
| (n of 30 scored video windows) | LD1 | 0.00 | (0.00, 0.50) | 0.00 | (0.00, 0.00) | .8574 | |
| | LD2 | 2.00 | (0.00, 7.00) | 0.00 | (0.00, 0.75) | | |
| | LD3 | 5.00 | (0.75, 14.50) | 0.00 | (0.00, 0.75) | | |
| | LD4 | 11.50 | (7.00, 14.25) | 0.00 | (0.00,6.75) | .0135 | * |
| Arched-back hovering/visible nursing | | | | | | | |
| (n of 30 scored video windows) | LD1 | 0.00 | (0.00, 0.00) | 0.00 | (0.00, 0.00) | 1.000 | |
| | LD2 | 0.50 | (0.00, 1.00) | 0.00 | (0.00, 0.00) | | |
| | LD3 | 1.50 | (0.00, 2.50) | 0.00 | (0.00, 0.00) | | |
| | LD4 | 2.00 | (1.00, 5.00) | 0.00 | (0.00, 1.75) | .0520 | |
| Pup interaction | | | | | | | |
| (n of 30 scored video windows) | LD1 | 12.5 | (8.75, 15.25) | 16 | (13.00, 18.50) | .1202 | |
| | LD2 | 9.5 | (8.00, 13.00) | 12.5 | (10.25, 16.00) | | |
| | LD3 | 10.5 | (7.75, 12.25) | 9 | (7.00, 14.25) | | |
| | LD4 | 10 | (7.75, 12.00) | 12.5 | (11.00, 16.25) | .0910 | |
| Nest building | | | | | | | |
| (n of 30 scored video windows) | LD1 | 19 | (12.25, 20.25) | 6.5 | (2.25, 12.25) | .0343 | * |
| | LD2 | 20 | (16.00, 25.25) | 11 | (4.25, 13.50) | | |
| | LD3 | 19 | (13.75, 23.50) | 10 | (5.00, 18.50) | | |
| | LD4 | 16.5 | (14.75, 20.25) | 9.5 | (4.25, 15.50) | .0439 | * |
| Solo activity | | | | | | | |
| (n of 30 scored video windows) | LD1 | 21 | (17.25, 24.50) | 28 | (17.75, 30.00) | .1042 | |
| | LD2 | 15 | (11.75, 19.25) | 24.5 | (23.25, 26.50) | | |
| | LD3 | 16.5 | (10.00, 22.50) | 24.5 | (14.75, 29.00) | | |
| | LD4 | 14.5 | (11.75, 19.25) | 24.5 | (16.00, 28.75) | .0206 | * |
| Time with pups | | | | | | | |
| (n of 30 scored video windows) | LD1 | 0.5 | (0.00, 3.75) | 0.5 | (0.00, 1.75) | .8599 | |
| | LD2 | 6.5 | (4.75, 9.50) | 0 | (0.00, 3.50) | | |
| | LD3 | 7 | (3.75, 15.25) | 0 | (0.00, 3.00) | | |
| | LD4 | 14 | (6.50, 16.50) | 6 | (0.00, 12.50) | .0508 | |
| Initiated pup retrieval | | | | | | | |
| (seconds from assay start, 0-1800) | LD1 | 1693.50 | (176.80, 1800) | 1292.50 | (163.80, 1800) | .8879 | |
| | LD2 | 225.5 | (15.25, 805.25) | 182 | (23.75, 1464.75) | | |
| | LD3 | 56.5 | (19.00, 479.20) | 1317 | (155.80, 1800) | | |
| | LD4 | 244.5 | (20.25, 513.50) | 576.5 | (219.80, 826.50) | .0697 | |
| Completed pup retrieval | | | | | | | |
| (seconds from assay start, 0-1800) | LD1 | 1800 | (1229, 1800) | 1800 | (1800, 1800) | .4017 | |
| | LD2 | 1158 | (962.80, 1365.20) | 1800 | (1385.00, 1800) | | |
| | LD3 | 898 | (755.20, 1011.50) | 1800 | (1286.00, 1800) | | |
| | LD4 | 658 | (612.00, 715.00) | 1564 | (1132.00, 1800) | .0000 | *** |
| Nest score | | | | | | | |
| (score from 0-4; | LD1 | 2 | (1.00, 3.00) | 0 | (0.00, 1.75) | .0752 | |
| 0, no nest attempted; | LD2 | 2.5 | (2.00, 3.25) | 1 | (0.25, 1.75) | | |
| 4, excellent nest) | LD3 | 3 | (2.00, 3.25) | 1 | (0.00, 3.50) | | |
| | LD4 | 3 | (3.00, 3.25) | 1.5 | (0.00, 4.00) | .2892 | |
