## Extended Figure 5-1 for "The lateral habenula is required for maternal behavior in the mouse dam"

| Variables and Group | Modeled term | Term value | CI 95% | df | *t* value | Pr (> l *t* l) | |
|---|---|---|---|---|---|---|---|
| All pups gathered | | | | | | | |
| Control dams | Intercept | -2.292 | (-6.368, 1.785) | 79.605 | -1.086 | 0.2809 | |
| DREADD dams | Intercept | 1.250 | (-4.796, 7.296) | 79.605 | 1.131 | 0.2613 | |
| Control dams | Slope | 5.025 | (3.680, 6.370) | 64.000 | 7.316 | 0.0000 | *** |
| DREADD dams | Slope | 1.060 | (-0.935, 3.055) | 64.000 | -3.892 | 0.0002 | *** |
| Nest building | | | | | | | |
| Control dams | Intercept | 16.583 | (12.416, 20.750) | 52.749 | 7.671 | 0.0000 | *** |
| DREADD dams | Intercept | 7.800 | (1.619, 13.980) | 52.749 | -2.739 | 0.0084 | ** |
| Control dams | Slope | 0.425 | (-0.691 1.541) | 64.000 | 0.746 | 0.4584 | |
| DREADD dams | Slope | 0.900 | (-0.755, 2.555) | 64.000 | 0.562 | 0.576 | |
