## Supplementary figures and images for "The lateral habenula is required for maternal behavior in the mouse dam"

### Extended Figure 6-1

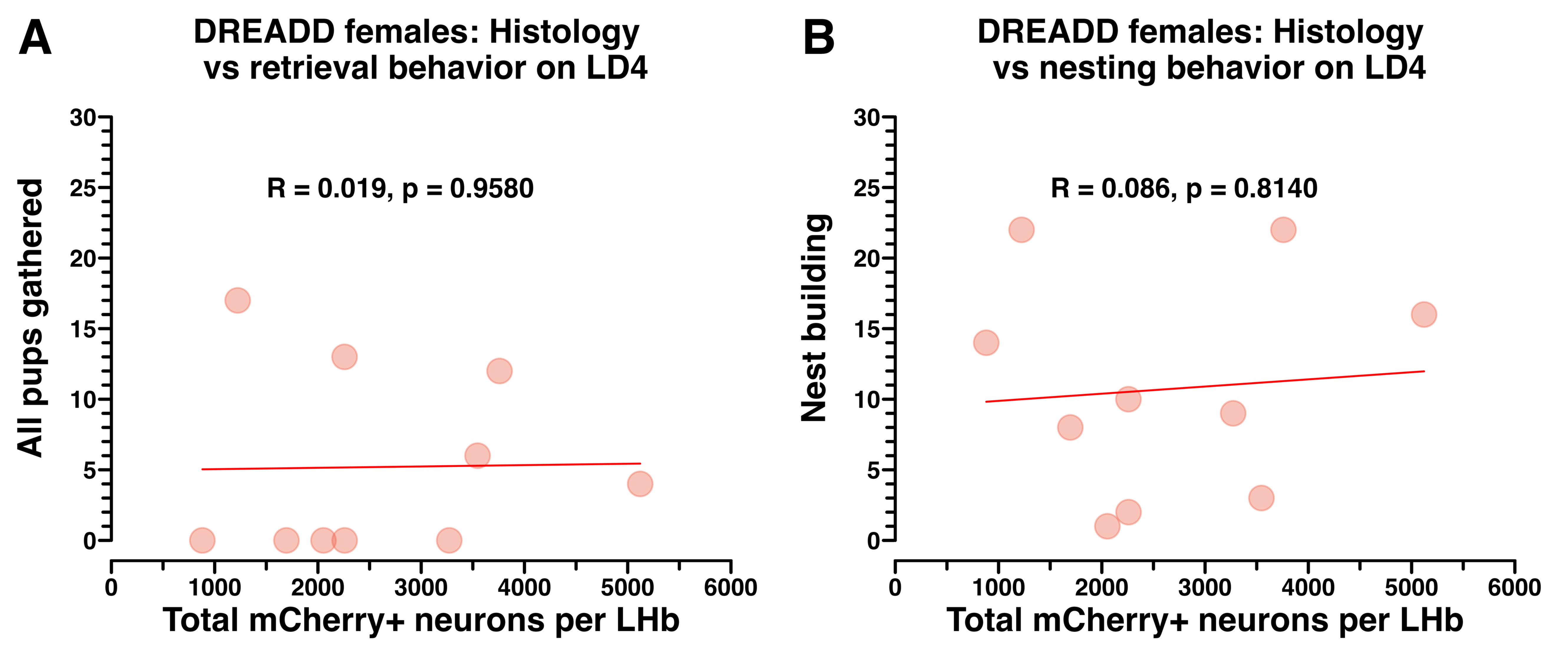

### Extended Figure 7-2

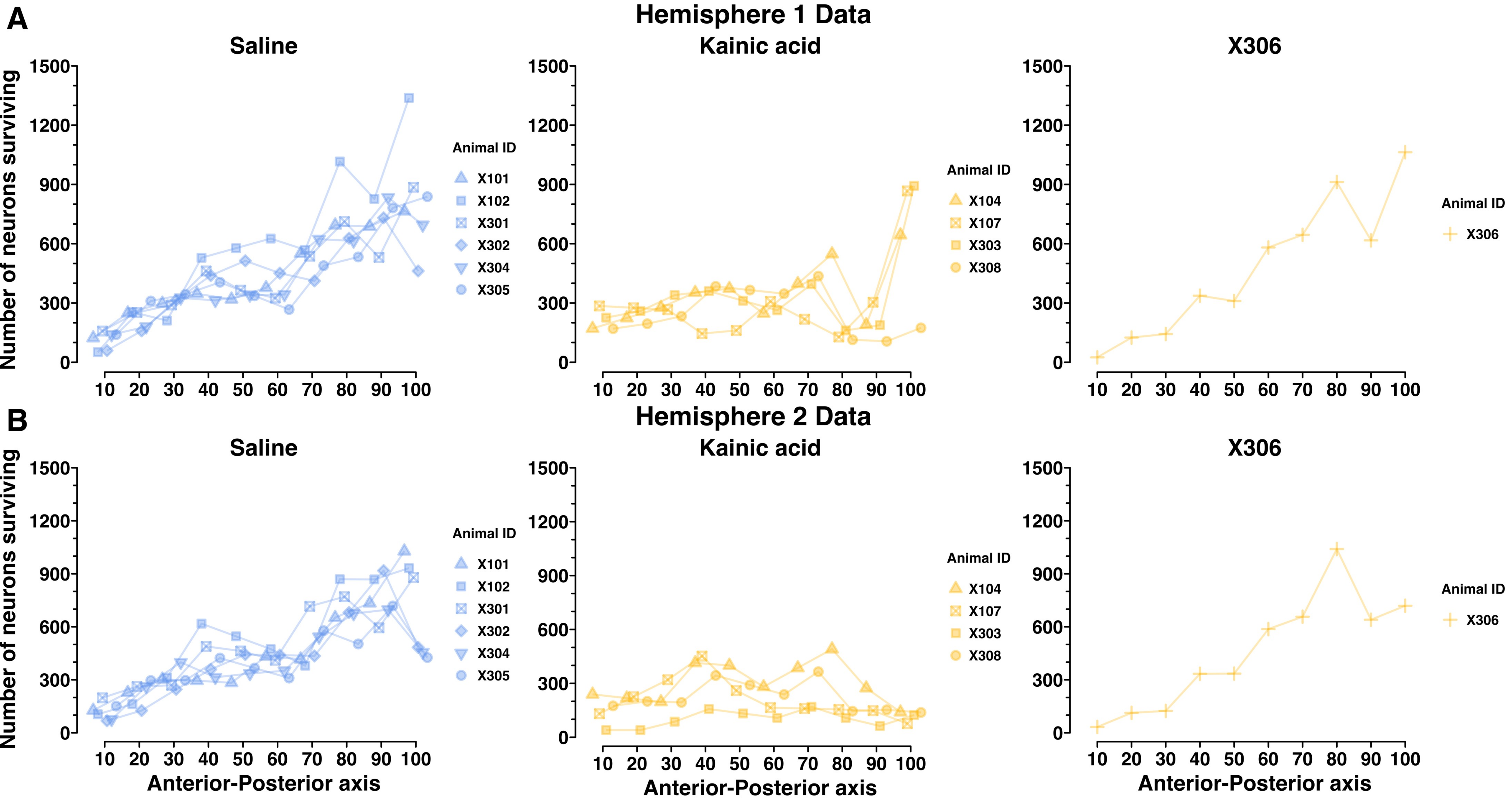
