## Extended Figure 6-2 for "The lateral habenula is required for maternal behavior in the mouse dam"

| Regions | Median Region scores DREADD dams | | All pups gathered *p* value | Nest building *p* value |
|---|---|---|---|---|
| Hippocampus, Median (IQR) | 2.05 | (1.70, 2.25) | 0.2777 | 0.2197 |
| Medial habenula, Median (IQR) | 0.2 | (0.10, 0.30) | 0.8704 | 0.5404 |
| Paraventricular nucleus of the thalamus, Median (IQR) | 0.55 | (0.300, 1.075) | 0.7321 | 0.2601 |
| Central lateral nucleus of the thalamus, Median (IQR) | 0.5 | (0.325, 0.775) | 0.179 | 0.2926 |
| Anterior pretectal nucleus, Median (IQR) | 3 | (2.00, 3.00) | 0.4147 | 0.8108 |
