## Extended Figure 7-1 for "The lateral habenula is required for maternal behavior in the mouse dam"

| Variables | Group | Modeled term | Term value | CI 95% | df | *t* value | Pr (> \| *t* \|) | |
|---|---|---|---|---|---|---|---|---|
| Hemisphere 1 | | | | | | | | |
| | Saline | 10th percentile | 130.367 | (59.489, 201.245) | 39.509 | 3.492 | 0.0012 | ** |
| | Kainic acid | 10th percentile | 211.316 | (99.248, 323.384) | 39.509 | 1.371 | 0.1780 | |
| | Saline | 100th percentile | 801.167 | (730.289, 872.045) | 39.509 | 21.459 | 0.0000 | *** |
| | Kainic acid | 100th percentile | 403.834 | (291.766, 515.902) | 39.509 | -6.731 | 0.0000 | *** |
| | Saline | Slope | 7.453 | (6.209, 8.698) | 88.000 | 11.734 | 0.0000 | *** |
| | Kainic acid | Slope | 2.139 | (0.171, 4.106) | 88.000 | -5.291 | 0.0000 | *** |
| Hemisphere 2 | | | | | | | | |
| | Saline | 10th percentile | 148.762 | (82.612, 214.912) | 23.074 | 4.260 | 0.0003 | *** |
| | Kainic acid | 10th percentile | 223.257 | (118.665, 327.849) | 23.074 | 1.349 | 0.1903 | |
| | Saline | 100th percentile | 757.721 | (691.571, 823.871) | 23.074 | 21.701 | 0.0000 | *** |
| | Kainic acid | 100th percentile | 196.993 | (92.401, 301.585) | 23.074 | -10.156 | 0.0000 | *** |
| | Saline | Slope | 6.766 | (5.775, 7.757) | 88.000 | 13.371 | 0.0000 | *** |
| | Kainic acid | Slope | -0.292 | (-1.859, 1.276) | 88.000 | -8.821 | 0.0000 | *** |
